# Dissociable neuronal substrates of visual feature attention and working memory

**DOI:** 10.1101/2023.03.01.530719

**Authors:** Diego Mendoza-Halliday, Haoran Xu, Frederico A.C. Azevedo, Robert Desimone

**Affiliations:** McGovern Institute for Brain Research, Massachusetts Institute of Technology, Cambridge, MA, USA

## Abstract

Attention and working memory (WM) are distinct cognitive functions, yet given their close interactions, it has been proposed that they share the same neuronal mechanisms. Here we show that in macaques performing a WM-guided feature attention task, the activity of most neurons in areas middle temporal (MT), medial superior temporal (MST), lateral intraparietal (LIP), and posterior lateral prefrontal cortex (LPFC-p) displays either WM coding or attentional modulation, but not both. One area thought to play a role in both functions is LPFC-p. To test this, we optogenetically inactivated LPFC-p bilaterally during the attention or WM periods of the task. Attention period inactivation reduced attentional modulation in LPFC-p, MST, and LIP neurons, and impaired task performance. WM period inactivation did not affect attentional modulation nor performance, and minimally reduced WM coding. Our results suggest that feature attention and WM have dissociable neuronal substrates, and that LPFC-p plays a critical role in attention but not WM.

## Introduction

Top-down visual attention and working memory (WM) are among the most studied cognitive functions and the most affected by neurological and psychiatric disorders. Visual WM allows us to temporarily store representations of stimuli when they become visually unavailable, whereas visual attention allows us to select, among all visually-available stimuli, those that are behaviorally relevant, and enhance their processing. Thus, while visual attention is most essential when relevant stimuli are visually present, visual WM is most essential when the relevant stimuli being memorized are visually absent.

Studies in humans using functional magnetic resonance imaging (fMRI) and electroencephalography (EEG) have reported a large overlap between the brain regions that play a role in WM and attention, most commonly including the lateral prefrontal cortex (LPFC) and parietal regions (Jonikaitis & Moore, 2019; Olivers, 2008). This anatomical overlap has led many researchers to claim that WM and attention share the same underlying neuronal mechanisms (Awh & Jonides, 2001), and to suggest that they are two constructs representing the same function (Olivers, 2008). However, most evidence for overlapping brain activity during WM and attention has come from studies in humans using fMRI and EEG methods in which signals reflect the aggregate activity of millions of neurons. Most single-neuron electrophysiological studies in non-human primates have examined either how the activity of neurons selectively encodes visual features maintained in WM (Konecky et al., 2017; Lara & Wallis, 2014; Leavitt et al., 2017; Mendoza-Halliday et al., 2014; Warden & Miller, 2010) or how neurons’ visual responses are modulated by attention to those features (Bichot et al., 2005; Ibos & Freedman, 2016; Maunsell & Treue, 2006; McAdams & Maunsell, 1999; Treue & Martínez Trujillo, 1999). Therefore, much less is known about whether the same or different neurons participate in these two mechanisms.

Another argument to support the claim that attention and WM share the same mechanisms is that they are strongly interdependent functions. Attention is thought to act as a gate-keeper for WM (i.e., attended stimuli are better encoded in WM; Awh & Jonides, 2001). In turn, it is believed that WM plays a key role in attention: attending to a visual stimulus feature is thought to require maintaining a WM representation of the target feature to be attended; this representation serves as an attentional template signal that, through top-down mechanisms, selectively modulates the activity of feature-tuned visual cortical neurons. One area that has been proposed as main source of top-down feature-attentional signals is LPFC (Desimone & Duncan, 1995; Gregoriou et al., 2014; Rossi et al., 2009), specifically the posterior subregion (LFPC-p; Bichot et al., 2015, 2019). Importantly, it has also been shown that LPFC-p neurons encode visual features maintained in WM (Mendoza-Halliday & Martinez-Trujillo, 2017). These findings raise two important questions: First, whether WM and attention signals are present in the same or different LPFC-p neurons; second, whether LPFC-p plays a critical role in both feature attention and WM maintenance.

The aim of this study was to investigate whether feature attention and WM share the same neuronal substrates, or whether they are dissociable. To examine this, we trained monkeys to perform a WM-guided spatially-global feature attention task for motion direction and recorded spiking activity from motion direction-selective neurons in multiple cortical areas at different processing stages: early visual area middle temporal (MT), visual association area medial superior temporal (MST), posterior parietal cortex (lateral intraparietal, LIP, 570 neurons), and LPFC-p. To test the causal role of LPFC-p in feature attention and WM, we optogenetically inactivated it bilaterally during the WM maintenance or sustained attention periods of the task and measured the effects on task performance and on the strength of WM and feature attention signals across the recorded areas.

## Results

We trained two rhesus macaque monkeys (*macaca mulatta*) to perform a WM-guided feature attention task (Fig. 1A). In each trial, a cue stimulus was presented for 0.8 s consisting of a full-screen random dot surface with coherent motion in one of two opposite directions. After a 3.2 s delay period during which monkeys maintained the cued direction in working memory (WM), a test stimulus was presented consisting of two overlapping full-screen random dot surfaces, one moving in the cued direction and the other in the opposite direction. Monkeys were trained to selectively attend to the surface with the cued direction (i.e., target surface) in order to detect and report (by releasing a hand-held lever) the occurrence of a small patch of dots with higher speed in the target surface while ignoring any such patch in the other surface (i.e., distractor surface). Because each changing patch was randomly located anywhere across the screen, monkeys were unable to predict the patch locations and were required to attend to all spatial locations. This design allowed our task to engage spatially-global feature attention and minimized the involvement of spatial attention during the test period before presentation of the target or distractor.

**Figure 1.**
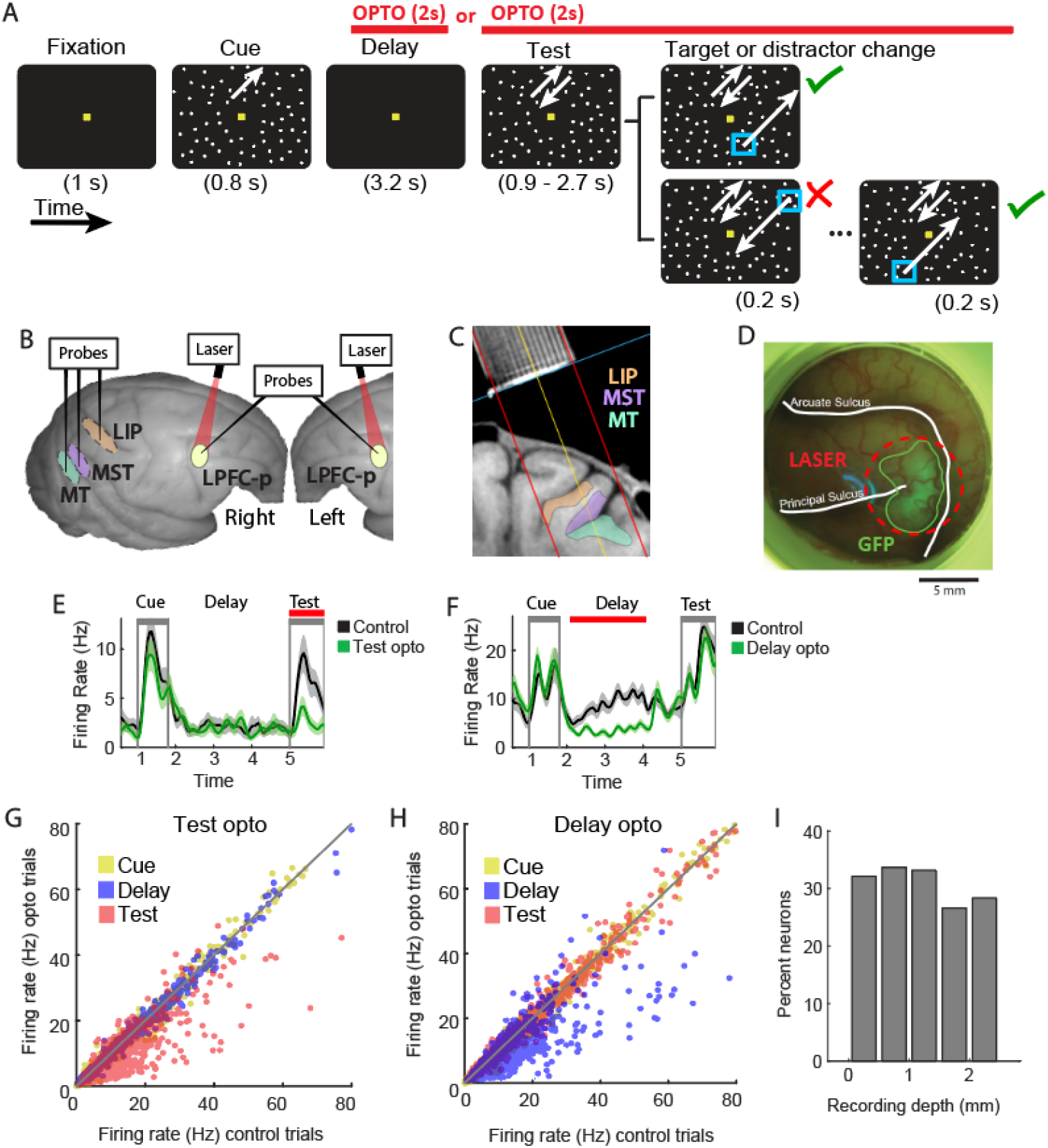
Behavioral, electrophysiological, and optogenetic methods. (A) Temporal sequence of behavioral task. White arrows depict motion directions of random dot surfaces; cyan squares depict speed-changing dot patches; red segments show periods of optogenetic LPFC-p inactivation; check mark and cross, correct and incorrect response periods, respectively. (B) Macaque brain photograph showing approximate location of cortical areas where electrophysiological probe recordings and optogenetic stimulation were performed. MT, MST and LIP are embedded within sulci. (C) Structural MRI nearly-coronal section of one monkey showing recording chamber grid (top) and location of areas LIP, MST and MT (colored) on the right hemisphere. Red lines show the orientation and range of available probe trajectories. (D) Photograph of an example LPFC chamber from one monkey showing cortical surface through transparent artificial dura, and region of viral expression indicated by GFP epifluorescence (enclosed by a green line). Dashed red line, area of laser illumination; white lines show sulci. Reflections of the light source on the artificial dura appear blue. (E,F) Mean firing rate (± standard error) of two example neurons over time across trials with (green) and without (black) optogenetic stimulation in the test period (E) or delay period (F). (G,H) Mean firing rate during the cue, delay and test periods across control trials (horizontal axis) and optogenetic inactivation (opto) trials (vertical axis) for all LPFC-p neurons recorded in test opto sessions (G) and delay opto sessions (H). (I) Firing rates of LPFC-p neurons before, during (red region) and after optogenetic stimulation, as a function of recording channel cortical depth.

In each monkey, we implanted 5 acute laminar probes in each experimental session to simultaneously record the spiking activity of motion direction-selective neurons from five cortical areas across a variety of processing stages: MT (938 neurons), MST (1865 neurons), LIP (570 neurons), and bilateral LPFC-p (1271 neurons), including posterior portions of areas 8Ad/v, 9/46d/v, and 45 (Fig. 1B,C; Paxinos, 2009). Because we simultaneously recorded multiple tens of neurons from a variety of brain areas and cortical depths, neurons were expected to vary widely in their receptive field locations and size; the spatially-global visual stimuli used in our task ensured that most recorded neurons were visually stimulated and engaged in the task independently of their receptive field location or size.

### Strength of working memory coding and feature attentional modulation across the visual processing stream

We first examined whether the delay period spiking activity of each single neuron encoded the cue direction held in WM, and whether its response to the test stimuli was modulated by the attended cue direction (Fig. 2A). We also estimated each neuron’s sensory selectivity for motion direction during the cue presentation period. Importantly, because there was no stimulus present during the delay period other than the fixation point, any differences in firing rates were indicative of selectivity for different memorized cue directions. Similarly, during the test period, we presented identical stimuli with the same combination of overlapping directions in all conditions with opposite cue directions (Fig. 1A), so any differences in test period firing rates between these conditions were due to cue direction-dependent attentional modulatory effects. We computed the area under the ROC curve (auROC, rectified to above 0.5) to estimate the discriminability between the distributions of firing rates in trials with opposite cue directions during the cue period (sensory discriminability), delay period (WM discriminability), and test period before target or distractor changes (attentional discriminability). These discriminability values served as a measure of the strength of sensory coding of motion direction, WM coding of motion direction, and feature attentional modulation, respectively. The presence or absence of discriminability was indicated by the statistical significance of the auROC value (see Methods).

**Figure 2.**
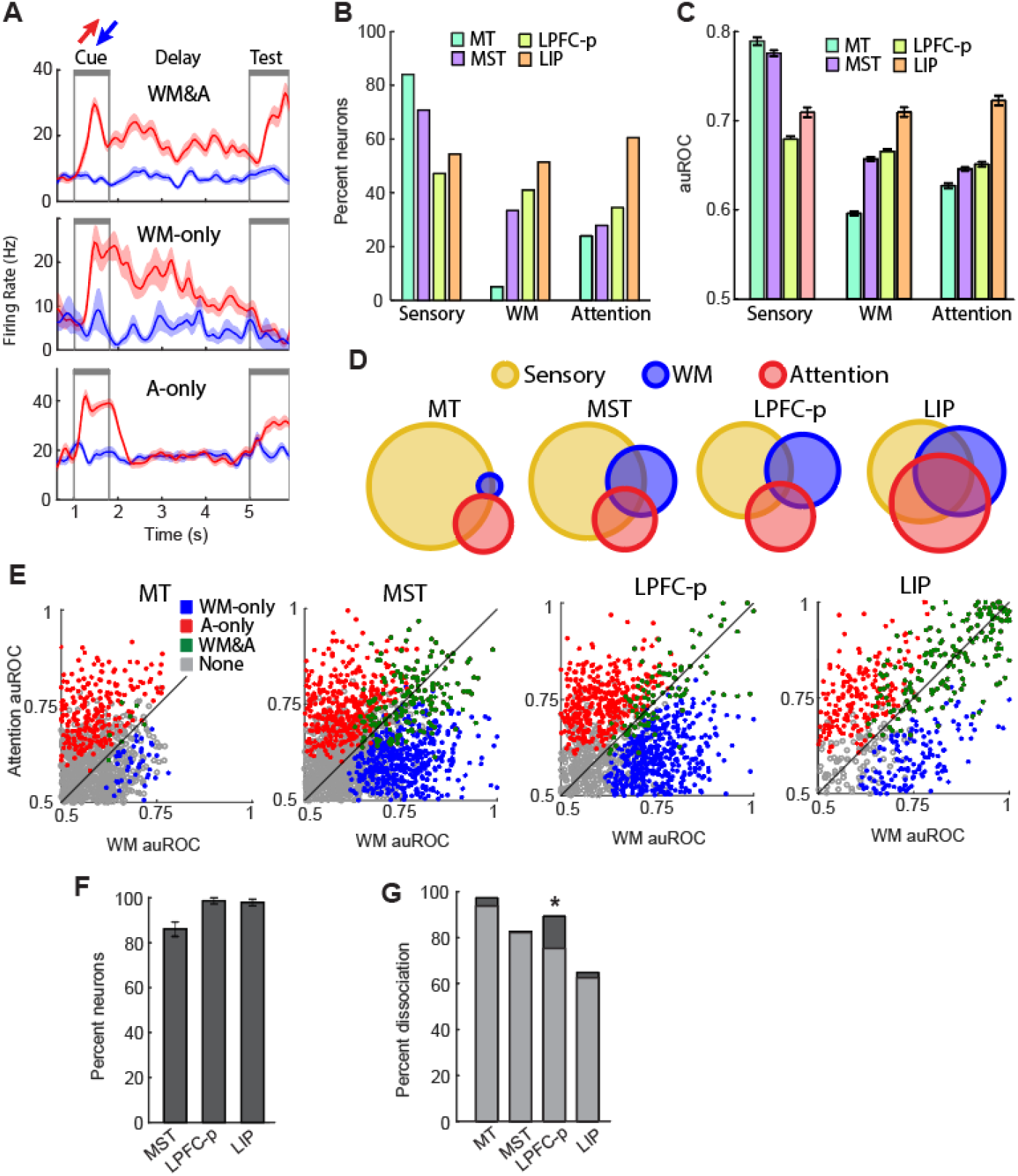
WM coding and feature attentional modulation in single neurons. (A) Mean firing rate (± standard error) over time across trials with opposite cue directions (red, blue) for three example neurons with different functional profiles. Top: Example neuron (LIP) showing WM coding and attentional modulation (WM&A). Middle: Example neuron (LIP) showing WM coding but not attentional modulation (WM&A). Bottom: Example neuron (MST) showing attentional modulation but not WM coding (WM&A). Gray lines, cue and test periods. (B) Percentage of neurons with significant sensory, WM, and attentional discriminability in each brain area. (C) Mean (± standard error) sensory, WM, and attentional discriminability across all neurons in each brain area. (D) Venn diagrams showing the relative proportion of neurons with significant sensory, WM, and/or attentional discriminability. Neurons with overlapping functions are represented by overlapping regions. (E) WM and attentional discriminability (auROC) of all individual neurons (dots). Neurons are classified (color-coded) by their significance of discriminability. (F) Percentage of neurons in each area for which WM coding and attentional modulation showed the same motion direction preference. (G) Among neurons with WM and/or attentional signals, percentage of neurons showing a dissociation between WM and attention (“WM-only” & “attention-only”). Overlaid light gray bars show the percent dissociation expected by random assignment (see Methods). *, significantly higher dissociation than expected by random assignment.

Next, we examined how sensory coding, WM coding and attentional modulation evolve across the visual processing stream, we compared sensory, WM, and attentional discriminability between the recorded areas. We found that overall, the percentage of neurons with significant sensory discriminability, as well as the mean sensory discriminability across all neurons, were highest in early visual cortex (MT) and followed a decreasing trend downstream (Fig. 2B-D). In contrast, for WM, the percentage of significant neurons, as well as and mean discriminability were particularly low in early visual cortex and increased progressively downstream (Fig. 2B-D). Similarly, for attention, the percentage of significant neurons and mean discriminability also increased in a downstream progression (Fig. 2B-D). These results were similar for both monkeys (Fig. S2), as were all subsequent results. The remaining figures show the results obtained from the combined neuronal populations from both monkeys.

### Single neuron-level dissociation between working memory and attentional modulation

Next, we examined whether WM and feature attention-related signals co-exist and interact within the same neurons, or whether these signals are dissociated among separate populations of neurons. We also asked whether the interaction of WM and attention signals differed across processing stages. If WM and attention are inseparable functions and share the same neuronal substrates, we would expect that neurons whose activity encoded the memorized directions would also show activity modulated by attention to these directions, and vice versa. Among all recorded neurons, we found that a fraction of them showed significant WM coding and significant feature attentional modulation (“WM & attention” neurons, Fig. 2A, top). However, a larger fraction of neurons showed either significant attentional modulation but not WM coding (“attention-only” neurons, Fig. 2A, center) or significant WM coding but not attentional modulation (“WM-only” neurons, Fig. 2A, bottom).

For each brain area, we classified all neurons into three groups – “WM & attention”, “WM-only”, and “attention-only” – based on whether their activity showed significant WM and/or attentional discriminability. We then obtained the percentage of neurons with each of these three functional profiles (Fig. 2D,E). Surprisingly, in all areas, the majority of neurons were “WM-only” or “attention-only”, and only a minority were “WM & attention” neurons. These results show that in all visual processing stages examined, WM and attention signals are, for the most part, segregated into separate populations of neurons, and that these signals co-exist or overlap only in a minority of neurons, between 3% in MT and 35% in LIP. Interestingly, in this minority, motion direction preference was the same during WM maintenance and attentional modulation (Fig. 2F), suggesting that feature preference in these neurons is a preserved property across the two cognitive functions.

We further found that the distribution of WM and attentional discriminability across the neuronal population differed widely between brain areas. First, while both WM and attentional discriminability were robustly present in MST, LPFC-p and LIP, we observed robust attentional discriminability but only weak and infrequent WM discriminability in MT, and WM and attentional discriminability were almost completely dissociated within neurons (Fig. 2E,G). Among the other three areas, the percentage of neurons showing a dissociation between WM and attentional discriminability was highest in LPFC-p and lowest in LIP (Fig. 2G). In LPFC-p, but not in other areas, this percentage was significantly higher than expected if WM and attentional discriminability were randomly and independently distributed between neurons given their observed incidences (see Methods). These results suggest that in all the brain areas examined, the substrates of feature attention and WM are mostly dissociated at the level of individual neurons, as well as at the brain area level – in MT.

### Large-scale bilateral optogenetic inactivation of LPFC-p

It has been proposed that LPFC-p plays a role in WM maintenance and as a source of top-down feature attention signals to other cortical areas. The presence of LPFC-p neurons showing WM coding and neurons showing attentional effects is compatible with this view, but it is not sufficient to determine whether the activity of these neurons plays a causal role in WM maintenance and feature attention. To test these causal roles, we developed a method for large-scale bilateral optogenetic LPFC-p inactivation during either the delay period (WM maintenance), or the test period (sustained feature attention). We implanted transparent artificial duras in the two cranial chambers covering left and right LPFC-p, adapting methods previously demonstrated by others (A. S. Nandy et al., 2017; Ruiz et al., 2013). The self-sealing property of the duras allowed us to perform approximately 170 injections of an AAV construct in 24 penetrations (12 per hemisphere) to drive neuronal expression of the red-shifted inhibitory opsin Jaws (Acker et al., 2016; Chuong et al., 2014) in a large area of left and right LPFC-p. Expression was confirmed *in vivo* by direct visualization of GFP epifluorescence on the cortical surface through the transparent artificial dura (Fig. 1D) starting two weeks after injections and before each recording session. Visualization also allowed us to estimate the average cortical surface area showing expression to be approximately 50 mm^2^ per hemisphere.

During each recording session, the right and left LPFC-p were simultaneously inactivated via external illumination of the cortical surface with red lasers placed 18 to 29 mm above the artificial duras. Our external illumination method allowed us to ensure optical stimulation of the entire region of opsin expression in each hemisphere (Fig. 1D). In some experimental sessions (“test-opto”, 22 in monkey Sh, 44 in monkey St), we optogenetically inactivated LPFC-p during the test period of the task, starting at the onset of the test presentation, and lasting 2s or until the monkey’s response. In other sessions, (“delay-opto”, 30 in monkey Sh, 39 in monkey St), inactivation was performed for 2s during the delay period starting 0.3s after cue offset and ending 0.9s before test onset. In both test-opto and delay-opto sessions, optical stimulation was randomly applied in half of the trials (i.e., opto trials), with the remaining trials serving as control (Fig. 1A).

In each experimental session, a recording probe was lowered in each prefrontal chamber at a random location within and around the region with visible opsin expression, allowing us to sample the activity of LPFC-p neurons during optogenetic stimulation. We confirmed that optogenetic stimulation during the test period caused decreases in the firing rates of LPFC-p neurons specifically during the test period (Fig. 1E, 1G 4A, S1A). Likewise, delay period optogenetic stimulation caused firing rate decreases that were specific to the delay period and did not extend into the test period (Fig. 1F, 1H, 4E, S1E). Across recording sessions, optogenetic modulation had a significant effect on firing rates in approximately 1/3 of the neurons. Of those, 93.5% showed a significant reduction in firing rates and 6.5% showed a significant increase. The percentage of neurons with significantly modulated activity was not very different across recording depths, although there was a somewhat lower percentage at deeper recording locations (Fig. 1I), likely due to decreases in light penetration of the cortex (Gong et al., 2020, Acker et al., 2016). Based on estimates of neuronal density (Collins et al., 2010; Young et al., 2013), on the percentage of modulated neurons, and on the mean surface area of the optically-stimulated expression region in both hemispheres, we estimated that on average across all recording sessions, the total number of LPFC-p neurons with significant response modulation in each session was approximately 5.1 million.

### LPFC-p inactivation during sustained feature attention impaired task performance

We first investigated whether optogenetic LPFC-p inactivation during the sustained feature attention period of the task (i.e., test period) impaired task performance. We found that the percentage of correct trials was lower in test opto trials than control trials in 100% of the 22 recording sessions for monkey Sh and in 93% of the 44 sessions for monkey St. Mean percent correct trials across sessions was significantly lower in opto trials than control trials for both monkeys (Fig. 3A; one-tailed paired t-tests, Sh: P < 0.0001, St: P < 0.0001). Moreover, mean reaction time across sessions was significantly higher in opto trials than in control trials for both monkeys (Fig. 3B; one-tailed paired t-tests, Sh: P = 0.0044, St: P = 0.0018). These results suggest that LFPC-p plays a causal role in feature attention.

**Figure 3.**
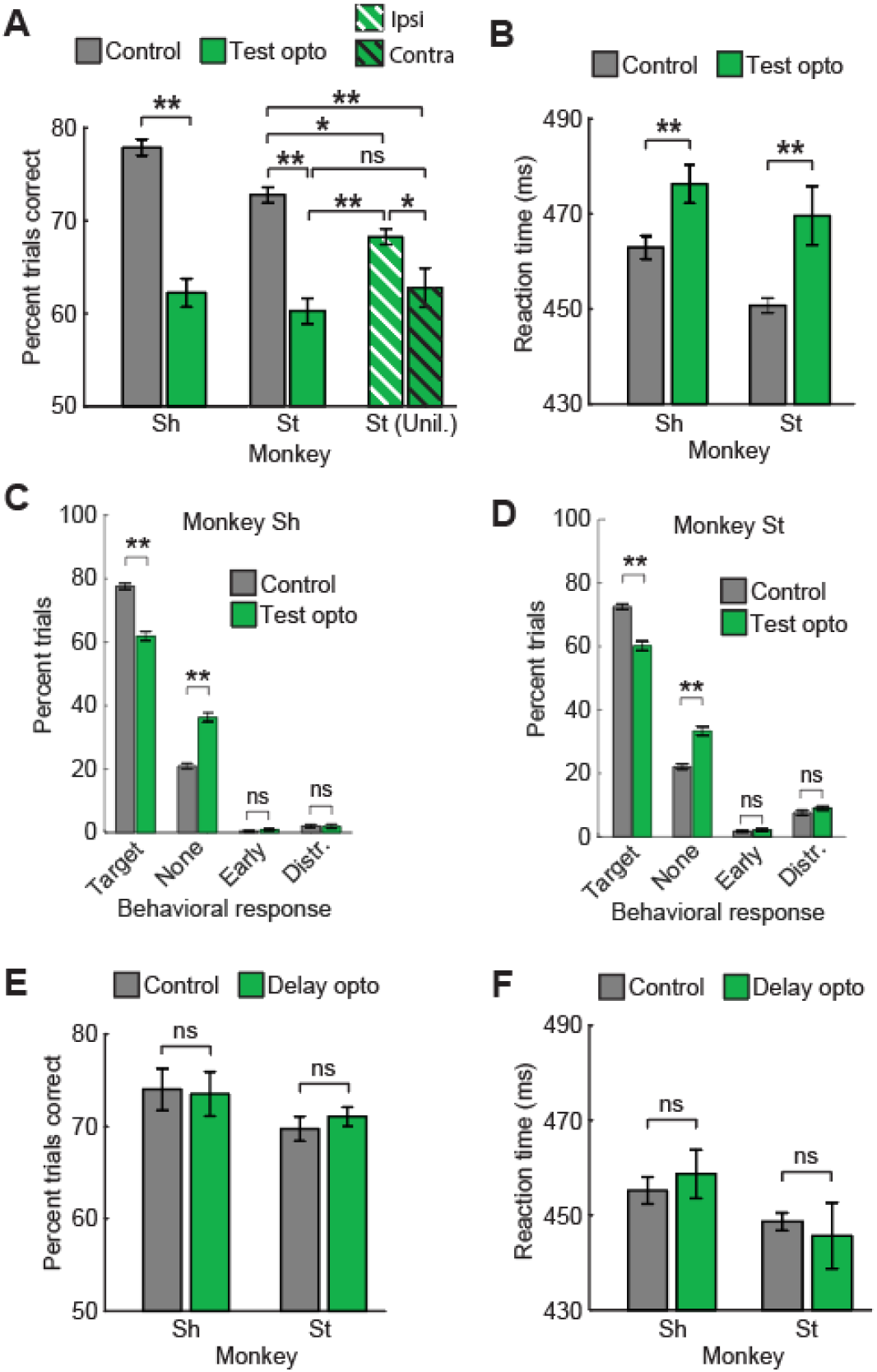
Behavioral effects of optogenetic prefrontal inactivation. (A,B) Mean (± standard error) task performance (A) and reaction time (B) across all test period optogenetic inactivation sessions in the control (gray) and opto (green) conditions, for monkeys Sh and St. Striped bars: mean task performance (A) and reaction time (B) for monkey St in unilateral prefrontal inactivation sessions, in trials with target changes in the hemifield ipsilateral (white stripes) or contralateral (black stripes) to the inactivated hemisphere. (C,D) Across-sessions mean (± standard error) percentage of trials with different types of behavioral responses, for monkeys Sh (C) and St (D). (E,F) Same as (A,B) for delay period optogenetic inactivation sessions. *, P < 0.05; **, P < 0.01; ns, not significant.

**Figure 4.**
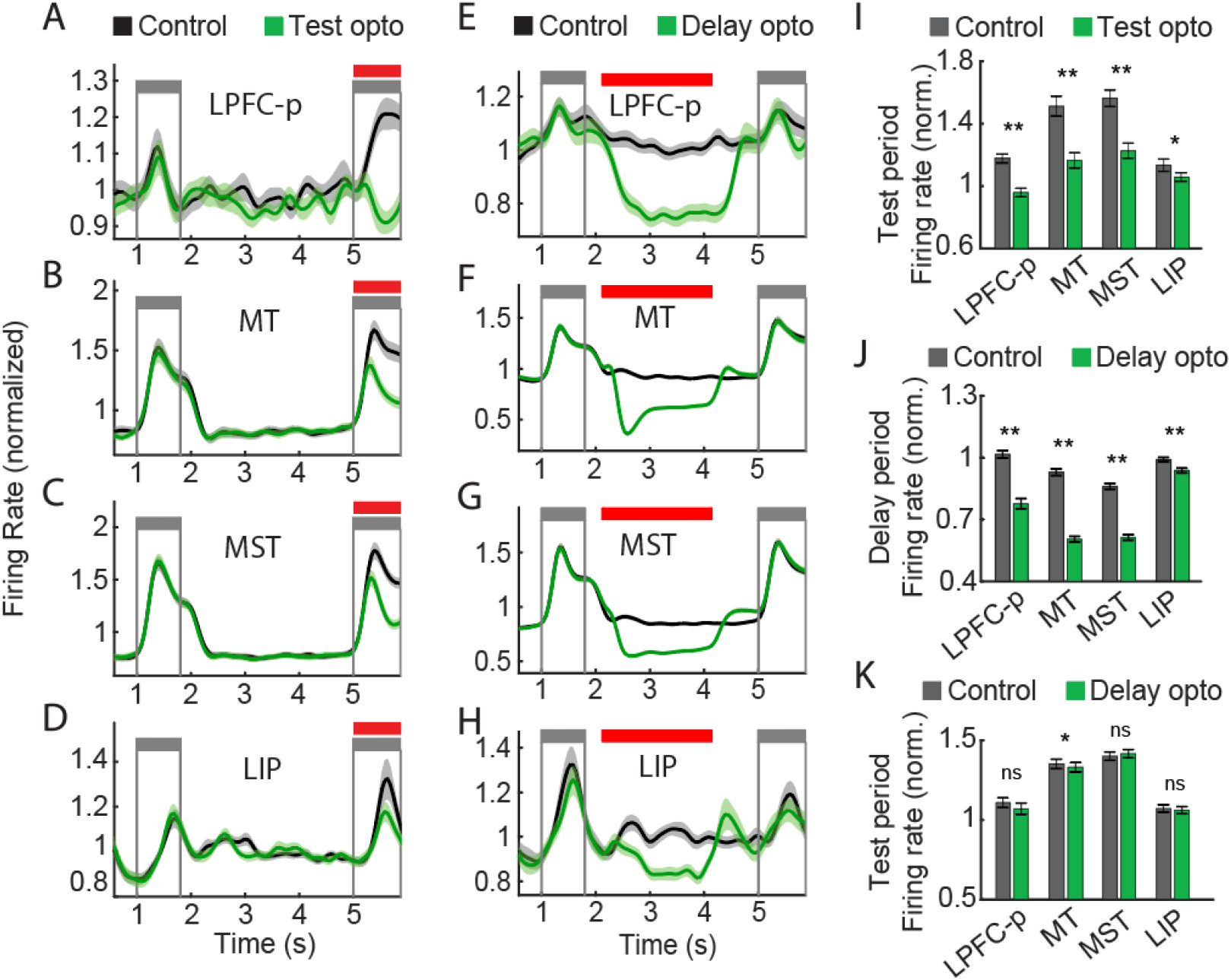
Effects of test and delay period optogenetic prefrontal inactivation on neuronal firing rates in each area. Mean normalized firing rate (± standard error) over time in control and opto trials across neurons with significant optogenetic modulation in LPFC, MT, MST, and LIP during sessions with optogenetic stimulation during the test period (A-D) or delay period (E-H). Gray lines, cue and test periods; red line, optogenetic stimulation period. (I) Mean test period normalized firing rate (± standard error) in control and opto trials across neurons in (A-D). (J-K) Mean delay period (J) and test period (K) normalized firing rate (± standard error) in control and opto trials across neurons in (E-H). *, P < 0.05; *, P < 0.01; ns, not significant.

To further examine the specificity of the behavioral impairments, we classified error trials into those with a response to the distractor change instead of the target, an early response (before the target, but not to the distractor), and no response. If LPFC-p inactivation caused a specific deficit in selective feature attention, it would impair detection of the targets (occurring in the attended dot surface) but not distractors (occurring in the unattended surface). Alternatively, if inactivation caused a non-selective perceptual deficit, it would impair detection of both targets and distractors. We tested these alternatives in Monkey St, which made a considerable fraction of distractor response errors. Despite inactivation causing a significant reduction in the percentage of target response trials (as reported above), it caused no significant reduction in the percentage of distractor response trials (P < 0.96; one-tailed paired *t-*test; Fig. 3D), suggesting that the impairment in task performance was in fact due to a deficit of selective feature attention. For both monkeys, inactivation caused no significant increase in the percentage of early error trials (Fig. 3C,D; one-tailed paired *t-*tests, Sh: P = 0.98; St: P = 0.25), indicating that the reduction in task performance was not due to an increase in response impulsivity caused by inactivation.

Previous studies have suggested that LPFC shows hemispheric lateralization during attention, with each cerebral hemisphere playing a stronger role when attention is allocated to the contralateral hemifield than the ipsilateral (Bichot et al., 2015; Buschman & Miller, 2007; Tremblay et al., 2015). To examine whether the behavioral effects of optogenetic LPFC-p inactivation reflect this lateralization, we carried out nine sessions in monkey St in which we inactivated LPFC-p unilaterally on the left or right hemisphere (rather than bilaterally) during the test period and compared task performance in trials in which the target speed change patch occurred on the visual hemifield ipsilateral or contralateral to the inactivated LPFC-p hemisphere. Consistent with the known functional lateralization, mean performance across sessions was significantly lower in the contralateral than the ipsilateral condition (Fig. 3A, one-tailed paired *t*-test, P = 0.023). Furthermore, mean performance across sessions in both unilateral inactivation conditions (ipsilateral and contralateral) was intermediate between that in the control condition and the bilateral inactivation condition (Fig. 3A). Therefore, unilateral LPFC-p inactivation is sufficient to impair task performance, but the impairment is more severe with bilateral than unilateral inactivation. The fact that the magnitude of the behavioral impairment is dependent on the relationship between the inactivated hemisphere and the visual hemifield of the target stimulus discards other potential non-specific effects of optogenetic stimulation as a cause of the observed behavioral impairments, such as increased response impulsivity or a motor impairment that would affect monkeys’ ability to release the hand-held lever to respond to the task.

### LPFC-p inactivation during the working memory delay did not impair task performance

We then examined whether bilateral LPFC-p inactivation during the delay period also had an effect on task performance. For both monkeys, there was no significant decrease in the mean percentage of correct trials (Fig. 3E; one-tailed paired t-tests, Sh: P = 0.33, St: P = 0.88) nor a significant increase in the mean reaction time (Fig. 3F; one-tailed paired t-tests, Sh: P = 0.26, St: P = 0.68) across sessions in the opto condition with respect to control. Together, our behavioral results show that bilateral optogenetic inactivation of LPFC-p during the test period leads to robust deficits in task performance, whereas the same inactivation during the delay period does not. These findings suggest that LPFC-p plays a critical role in feature attention but not in WM maintenance, and that these two functions have dissociable mechanisms in LPFC-p.

Based on these behavioral results, we reasoned that optogenetic LPFC-p stimulation during the test period caused alterations in neuronal activity that impaired performance of the task. Furthermore, given the known feedback projections from LPFC-p to visual and parietal areas that likely play a role in the task, including MT, MST and LIP, we further speculated that LPFC-p inactivation also led to changes in neuronal activity in these distant cortical areas; in contrast, we predicted that optogenetic LPFC-p stimulation during the delay period would modulate activity in a manner that minimally affects task performance.

### Test period LPFC-p inactivation decreased attentional modulation of neuronal responses

We investigated the effects of LPFC-p inactivation on the spiking activity of neurons in all the recorded areas. First, we compared the mean firing rates of neurons in each area across all opto trials and control trials. As expected, optogenetic stimulation during the test period drastically decreased the firing rates of neurons in LPFC-p (Fig. 4A,I, S1A). Interestingly, test period stimulation also led to a significant reduction in the firing rate of neurons in MT, MST, and LIP (to a lesser extent) – regions located far from the optogenetically stimulated region (Fig. 4B-D,I, S1B-D; paired *t*-tests). Thus, changes in LPFC-p activity have modulatory effects on the activity of visual and parietal neurons, possibly via feedback connections (Mendoza-Halliday et al., 2014; Xu et al., 2022).

Next, we examined whether optogenetic LPFC-p inactivation during the test period caused a reduction in the strength of feature attentional modulation in LPFC-p and in other recorded areas (Fig. 5A). As before, we measured the strength of attentional effects for each neuron using attentional discriminability, i.e., the auROC between the distribution of test period firing rates in trials with preferred vs. anti-preferred cue directions. This was done independently for control and opto trials. Among all neurons with optogenetically-modulated firing rates, LPFC-p inactivation significantly decreased the strength of attentional effects in LPFC-p (Fig. 5B,F), as well as in MST (Fig. 5D,H; Wilcoxon rank sum tests; see Methods). In these two areas, this effect was also significant across the entire population of neurons with attentional effects, including neurons with and without a significant optogenetic effect on overall firing rates (Fig. 5J; Wilcoxon rank sum tests; see Methods). Inactivation also reduced the percentage of LPFC-p and MST neurons with significant attentional effects (Fig. 5K). Although prefrontal inactivation caused an overall decrease in firing rates in MT and LIP neurons (Fig. 4B,D, S1B,D & 5C,E), it did not significantly reduce the mean attentional effect across neurons with optogenetically-modulated firing rates (Fig. 5G,I; Wilcoxon rank sum tests; see Methods) nor across all neurons with significant attentional effects (Fig. 5J,K; Wilcoxon rank sum tests; see Methods). Consistent with these results, the mean change in attentional discriminability by prefrontal inactivation was significantly larger in MST than MT and LIP (unpaired *t*-tests, P < 0.001).

**Figure 5.**
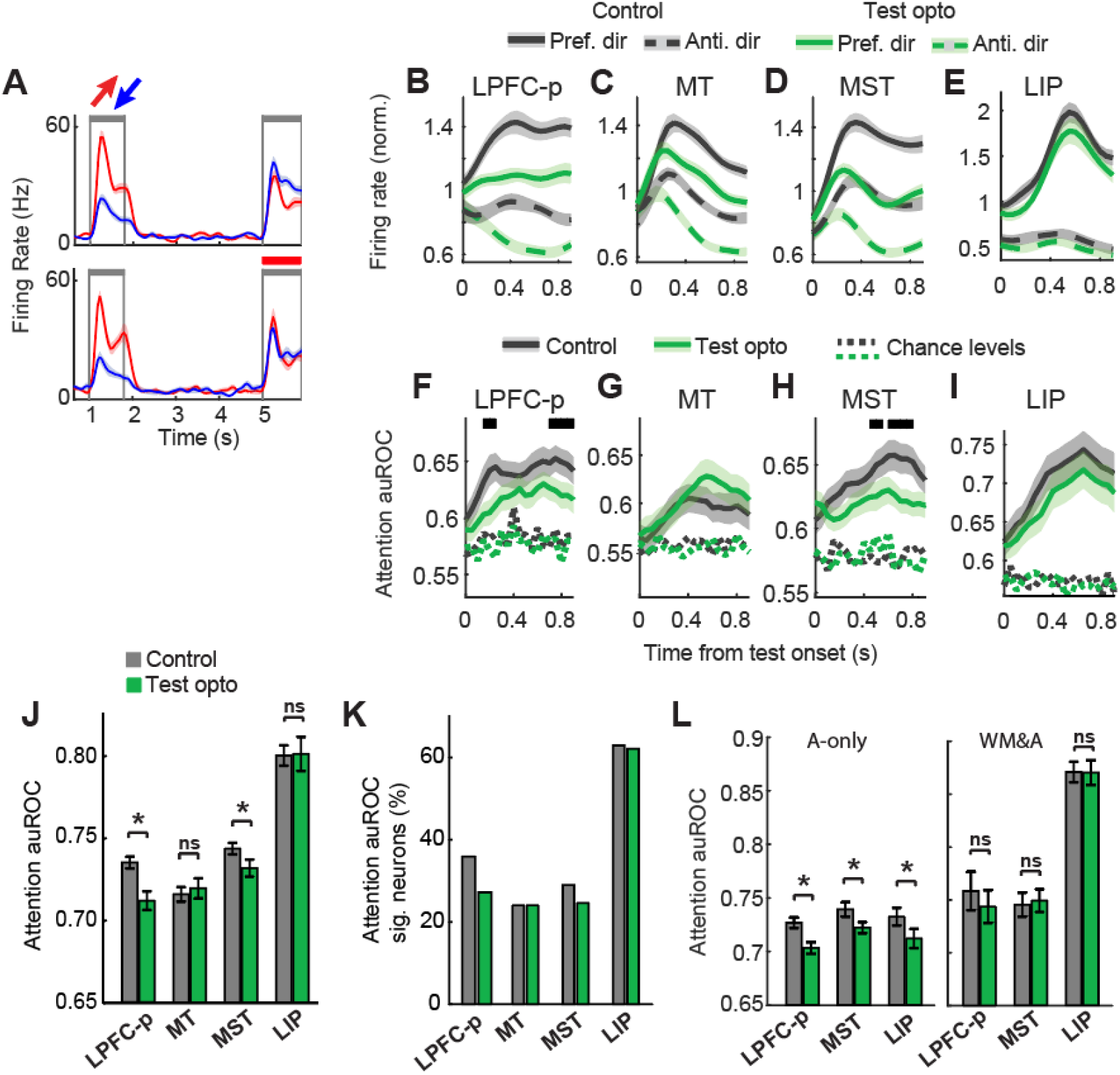
Effects of test period optogenetic prefrontal inactivation on attentional modulation. (A) Mean firing rate (± standard error) over time across trials with opposite cue directions (red, blue) for an example neuron (MST) with significantly lower attentional discriminability in test opto trials (bottom) than control trials (top). Gray lines, cue and test periods; red line, optogenetic stimulation period. (B-E) Mean normalized firing rate during the test period across neurons in each brain area with significant optogenetic firing rate modulation, in control and test opto trials with preferred and anti-preferred attended directions. (F-I) Mean auROC during the test period across neurons in each brain area with significant optogenetic firing rate modulation in control and test opto trials. Black segments, time bins with significant auROC difference between control and opto trials; dotted lines, mean auROC expected by chance in control and opto trials. (J) Mean test auROC (± standard error) across neurons in each brain area with significant attentional discriminability in control and test opto trials. (K) Percentage of neurons in each brain area with significant attentional discriminability in control and test opto trials. (L) Mean test auROC (± standard error) across “attention-only” neurons (left) and “WM & attention” neurons (right) –in each brain area in control and test opto trials. *, P < 0.05; ns, not significant.

As reported earlier, among neurons with attentional effects, some showed WM coding (“WM & attention” neurons) while others did not (“attention-only” neurons). We investigated whether attentional modulation strength was affected differently by LPFC-p inactivation in “WM & attention” neurons and in “attention-only” neurons. This comparison was possible in LPFC-p, MST, and LIP, where WM coding was considerable present in the neuronal population. In all three areas, inactivation during the test period led to a significant reduction in mean attentional discriminability among “attention-only” but not among “WM & attention” neurons (Fig. 5L; Wilcoxon rank sum tests; see Methods). Therefore, LPFC-p inactivation reduced the strength of attentional signals in these three areas, but only in neurons with no WM coding. The observation that LPFC-p inactivation reduced attentional effects not only locally but also distantly (in MST and LIP) supports the idea that LPFC-p serves as a source of feature attentional feedback signals that modulate neuronal responses in cortical areas along the visual processing stream. The reduction of attentional effects with test period LPFC-p inactivation may explain the observed impairments in task performance.

### Effects of delay period LPFC-p inactivation on working memory coding

The lack of behavioral effects from delay period optogenetic LPFC-p inactivation suggested that there was a fundamental difference in the neuronal effects of delay period optogenetic stimulation compared to test period stimulation. We first examined whether delay period optogenetic stimulation had an overall effect on neuronal firing rates. Optogenetic LPFC-p stimulation during the delay period decreased the firing rates of neurons in all recorded areas, both locally in LPFC-p as well as distantly in MT, MST and LIP (Fig. 4E-H,J; paired *t*-tests, all P < 0.01). Importantly, this effect was limited to the delay period, and did not extend to the test period (Fig. 4K). These results imply that the lack of effects of delay period optogenetic stimulation on task performance cannot be due to a failure of our optogenetic methods to reduce delay period neuronal activity in LPFC-p and other recorded areas; instead, it is possible that behavior was unaffected because delay period optogenetic stimulation did not modulate activity during the test period, when sustained feature attention was implemented.

Next, we examined whether optogenetic LPFC-p inactivation during the delay period affected the ability of neurons to encode the cue direction in WM, as measured by WM discriminability. We investigated this in LPFC-p, MST and LIP, where there was substantial WM discriminability in the neuronal population. While optogenetic stimulation caused an overall reduction in delay period firing rates in LPFC-p (Fig. 4E), this reduction was similar across cue direction conditions and did not lead to a significant reduction in WM discriminability (Fig. 6B,E,H,I; Wilcoxon rank sum tests; see Methods). In contrast, in MST, delay period optogenetic stimulation caused a significant reduction in WM discriminability among neurons with a delay period optogenetic modulation of firing rates (Fig. 6A,C,F Wilcoxon rank sum tests). This effect was also significant across all neurons with WM discriminability, including those with and without a significant optogenetic effect on overall firing rates (Fig. 6H; Wilcoxon rank sum test). Optogenetic stimulation also reduced the percentage of MST neurons with significant WM discriminability (Fig. 6I). However, the optogenetic effect on WM discriminability in MST was only temporary, and was not visible soon after the offset of optogenetic stimulation (Fig. 6F). In LIP, there was no significant effect on WM discriminability (Fig. 6D,G,H; Wilcoxon rank sum tests). In sum, LPFC-p inactivation led to a temporary reduction in the strength of WM coding in MST. In contrast, in LPFC-p and LIP, WM coding was robust to and unaffected by optogenetic modulations of firing rates.

**Figure 6.**
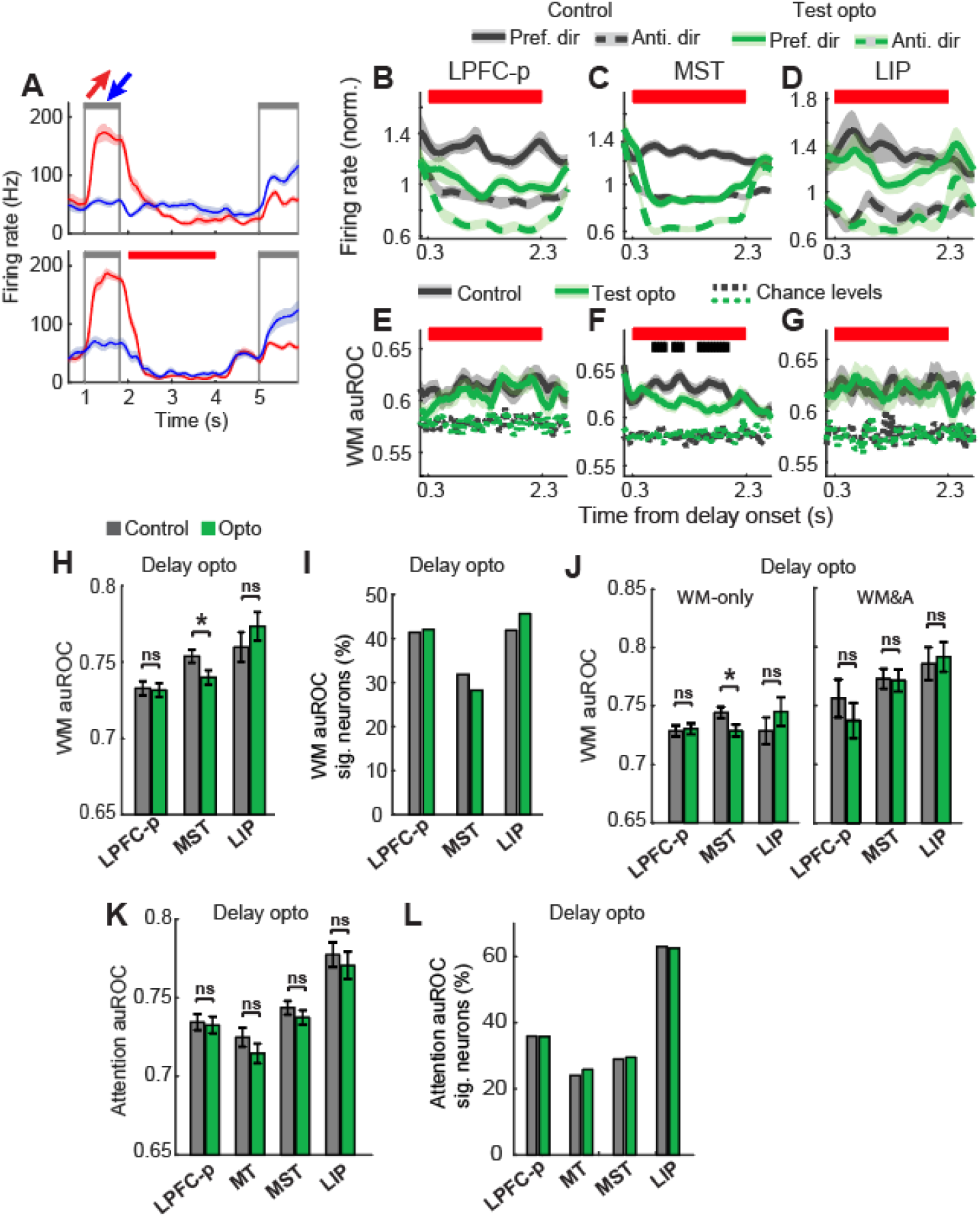
Effects of delay period optogenetic prefrontal inactivation on attentional modulation. (A) Mean firing rate (± standard error) over time across trials with opposite cue directions (red, blue) for an example neuron (MST) with significantly lower WM discriminability in delay period opto trials (bottom) than control trials (top). Gray lines, cue and test periods; red line, optogenetic stimulation period. (B-D) Mean normalized firing rate during the delay period across neurons in each brain area with significant optogenetic firing rate modulation, in control and test opto trials with preferred and anti-preferred memorized directions. (E-G) Mean auROC during the delay period across neurons in each brain area with significant optogenetic firing rate modulation, in control and delay opto trials. Red segment, optogenetic stimulation period; black segments, time bins with significant auROC difference between control and opto trials; dotted lines, mean auROC expected by chance in control and opto trials. (H) Mean delay auROC (± standard error) across neurons in each brain area with significant WM discriminability in control and delay opto trials. (I) Percentage of neurons in each brain area with significant WM discriminability in control and delay opto trials. *, P < 0.05; ns, not significant. (J) Mean delay auROC (± standard error) across “WM-only” neurons (left) and “WM & attention” neurons (right) in each brain area in control and delay opto trials. (K) Mean test auROC (± standard error) across neurons in each brain area with significant attentional discriminability in control and delay opto trials. (L) Percentage of neurons in each brain area with significant attentional discriminability in control and delay opto trials. *, P < 0.05; ns, not significant.

We then asked whether in areas with considerable WM and attentional signals (LPFC-p, LIP and MST), LPFC-p inactivation had different effects on “WM & attention” neurons and “WM-only” neurons. In MST, inactivation during the delay period significantly decreased mean WM discriminability among “WM-only” neurons but not “WM & attention” neurons (Fig. 6J; Wilcoxon rank sum tests). Thus, the effect of prefrontal inactivation on WM coding was limited to MST neurons that exclusively carry WM signals but not attentional signals. In LPFC-p and LIP, there was no significant effect on WM discriminability for any of these neuron types (Fig. 6J; Wilcoxon rank sum tests).

### Delay period LPFC-p inactivation did not reduce attentional modulation

The observation that optogenetic prefrontal inactivation decreased task performance when performed during the test period, but not during the delay period, suggests that test period inactivation affects neuronal activity that is critical to the feature attention task, whereas delay period inactivation does not. We speculated that the critical activity may be that which is modulated by attention during the test presentation and allows the preferential processing of target speed changes. We therefore examined whether delay period prefrontal inactivation modulates neuronal activity during the test period, and whether it has an effect on the strength of attentional modulation in the recorded areas.

We found that delay period optogenetic stimulation did not modulate overall firing rates during the test period in any of the recorded areas (Fig. 4E-H, K; paired *t*-tests, P ≥ 0.05; a significant but very small 1.5% firing rate reduction was observed in MT), despite causing significant decreases in firing rates during the delay period in all areas (Fig. 4J; paired *t*-tests, P < 0.05). Likewise, delay period optogenetic stimulation also did not result in significant decreases in the strength of attentional effects in any of the areas (Fig. 6K; Wilcoxon rank sum tests), nor decreases in the percentage of neurons with significant attentional effects (Fig. 6L). The presence of both a decrease in task performance and a decrease in attentional effect strength in LPFC-p and MST with test period prefrontal inactivation, and the absence of both of these effects with delay period inactivation, suggests that the observed behavioral deficit may be the result of decreases in attentional modulatory signals.

## Discussion

To examine whether the neuronal substrates underlying feature attention and working memory (WM) are dissociable, here we recorded the activity of neurons in multiple cortical visual processing stages of macaque monkeys performing a spatially-global WM-guided feature attention task for motion direction. To our knowledge, this is the first task in non-human primates using spatially-global stimuli and requiring feature attention to be allocated evenly across spatial locations, therefore minimizing the recruitment of spatial attention.

In all areas, we found neurons with delay period activity encoding the directions in WM and neurons with a response to the attended test stimuli that was modulated by the attended direction. This is in agreement with several human fMRI and EEG studies that have reported an overlap between brain regions with WM-related and feature attention-related activity (Jonikaitis & Moore, 2019; Olivers, 2008). However, we further found that the majority of neurons in all areas encoded motion directions in WM or were modulated by attention to motion directions, but not both. Our results strongly suggest that within each area, different neurons contribute to the mechanisms of feature attention and WM. Consistent with this finding, previous studies have shown that, at least in LPFC, spatial locations and memorized locations are encoded mostly by different neurons (Lebedev et al., 2004; Messinger et al., 2009). It is likely that the single neuron-level dissociation we report here was not detectable by the aforementioned human fMRI and EEG studies due to their low spatial resolution (Leavitt et al., 2017). Moreover, in agreement with previous studies, we found that in early visual area MT, feature attention strongly modulated the activity of many neurons (Treue & Martínez Trujillo, 1999), but WM signals were weak or absent (Mendoza-Halliday et al., 2014; Zaksas & Pasternak, 2006). This suggests that in early visual processing stages, neuronal activity is modulated by feature attention but not WM – an example of a region-level dissociation between these two functions. Interestingly, both WM coding and attentional modulation were much stronger in LIP than LPFC-p. Future studies inactivating LIP during similar tasks may further clarify whether LIP plays a more critical role in feature attention or WM maintenance than LPFC-p.

To examine whether LPFC-p plays a causal role in feature attention and WM, we performed bilateral optogenetic LPFC-p inactivation. Innumerable studies in small species such as mice and rats have successfully achieved robust behavioral effects by optogenetically perturbing neuronal activity in specific brain regions (Fenno et al., 2011). In contrast, a much lower fraction of macaque studies have successfully achieved optogenetically-induced behavioral effects, and these effects are often weak (Tremblay et al., 2020). One potential reason is that given the much larger brain size in macaques than mice and rats, the size of the perturbed regions using standard optogenetic approaches only represents a small fraction of typical functional regions in the macaque brain (Gong et al., 2020). In our study, we aimed to inactivate a large surface area in the posterior LPFC bilaterally. To accomplish this, we implemented multiple improvements to traditional approaches. By replacing the dura with a self-sealing transparent artificial dura (A. S. Nandy et al., 2017; Ruiz et al., 2013), we were able to perform a large number of virus injections with anatomical precision, daily visualization of epifluorescence to track the extent of opsin expression over time, electrode penetrations for neuronal recordings, and external laser stimulation of a large surface area customized to match the expression region. While various studies have performed optogenetic stimulation through an artificial dura (Nandy et al., 2019; Nassi et al., 2015), to our knowledge, none of these studies has performed optogenetic inactivation of such a large cortical surface bilaterally. Given the success of our study at achieving large-scale silencing of neuronal activity and robust behavioral effects, our methods provide a means for future studies to silence other large superficial cortical areas and measure the impact on behavior.

We bilaterally inactivated LPFC-p during the test period to examine its role in feature attention. In both monkeys, test period inactivation caused a strong impairment in task performance. This impairment was accounted for by a selective reduction in detection of targets but not distractors. These results suggest that LPFC-p plays a causal role in selective feature attention. We also found that LPFC-p inactivation reduced the strength of attentional effects on neuronal activity in LPFC-p, MST and LIP. This suggests that LPFC-p plays a critical role as a source of top-down feature attentional signals that selectively modulate stimulus responses in visual and parietal cortical neurons. Our results are consistent with prior studies showing that muscimol inactivation of the ventral prearcuate region within LPFC-p impaired performance of a feature search task and reduced the effects of feature attention in FEF and V4 neurons (Bichot et al., 2015). Interestingly, the observation that LPFC-p inactivation reduced attentional effects in “attention-only” neurons but not in “WM and attention” neurons in MST and LIP indicates that the top-down modulatory mechanisms of feature attention exclusively target neurons that are not involved in WM coding. This shows another dissociation between the neuronal substrates of feature attention and WM. Why we found no significant decrease in attentional effects in MT with LPFC-p inactivation remains unclear. One possibility is that the attentional modulatory influence of LPFC-p is stronger in areas closer along the visual hierarchy such as LIP and MST, and weaker and harder to detect in more distant areas such as MT.

One of the most remarkable findings in our study was that while task performance was strongly impaired by test period LPFC-p inactivation, it was unaffected by delay period inactivation. This behavioral dissociation suggests that LPFC-p plays a critical role in feature attention but not in WM maintenance. Consistent with this, we also found that delay period inactivation did not reduce the strength of WM representations in LPFC-p nor LIP neurons, and only did so in “WM-only” neurons in MST. This effect was confined to the inactivation period and did not get carried over to the test period, indicating that WM representations were restored in MST after the offset of inactivation.

The lack of behavioral effects of delay period LPFC-p inactivation suggests that the activity of LPFC-p neurons encoding WM representations observed in our study and others (Konecky et al., 2017; Lara & Wallis, 2014; Leavitt et al., 2017; Mendoza-Halliday et al., 2014; Warden & Miller, 2010) is not necessary for WM storage. There are several possible reasons why this is the case: first, lesion studies in monkeys have proposed that LPFC is not necessary for mere storage of WM representations, and that more anterior and dorsal LPFC subregions (excluding LPFC-p) play a critical role when WM representations are actively monitored or manipulated (Petrides, 1991, 2005). A second proposed alternative that remains mostly theoretical is that LPFC-p contributes to WM storage, but it does so via synaptic or other activity-silent mechanism – one that does not depend on spiking activity (Mongillo et al., 2008; Stokes, 2015). Lastly, it has been suggested that WM representations are stored in the spiking activity of a widely distributed network of brain regions, and that no one region plays a critical role in WM storage (Christophel et al., 2017). This is consistent with our finding that WM representations of motion direction are simultaneously encoded by neurons in at least three areas – MST, LIP, and LPFC-p (and perhaps more). In contrast to the distributed nature of WM storage, our findings suggest that the mechanisms of feature attentional modulation are less distributed and rely more critically on LPFC-p.

In summary, we showed that in MT, MST, LIP, and LPFC-p, the activity of most neurons displays either WM coding or attentional effects but not both. We also showed that LPFC-p plays a causal role in feature attentional modulation but not in WM. Dissociations between various neural and behavioral signatures of attention and WM have been reported by several studies using EEG (Bae & Luck, 2018; Harris et al., 2020), fMRI (Mayer et al., 2010; Tomasi et al., 2007), and behavioral measures (Tas et al., 2016) in healthy subjects, as well as in subjects with attention-deficit/hyperactivity disorder (ADHD; Mattfeld et al., 2016) and fronto-parietal strokes (Peers et al., 2020). Complementing these findings, our results indicate that feature attention and WM do not rely on the same neuronal substrates and suggest that they are two distinct functions rather than two constructs representing the same function.

## Methods

### Animal model and subjects

Two male Rhesus macaque monkeys (*macaca mulatta*) were used in this study: monkey Sh (9 years old and 13.7 kg) and monkey St (10 years old and 12.1 kg). Prior to this study, none of the animals had participated in another study nor been subject to surgical procedures. The animals were housed on 12-hr day/night cycles and maintained in a temperature-controlled environment (80°F). They were provided food and water *ad libitum* except during days in which they participated in behavioral training or experimental sessions. On these days, instead of water, they received apple juice as reward for correct task performance during the procedures at the lab. Training and experimental sessions were carried during afternoon hours, before 7pm. All procedures were approved by the MIT Committee on Animal Care and were in accordance with the guidelines of the National Institutes of Health Guide for the Care and Use of Laboratory Animals.

### Behavioral task and performance

The behavioral task was run on a PC computer using Psychomonkey/Psychtoolbox in Matlab (MathWorks, USA). Visual stimuli were presented on an LCD monitor (61 × 34.3 cm) with 120 Hz refresh rate (Acer, Taiwan). During task performance, animals sat on a plexiglass chair at a viewing distance of 57cm and were head-fixed using a headpost. Eye position was tracked using a camera-based Eyelink 2 system with 500 Hz sampling rate.

To examine the mechanisms underlying feature attention and working memory (WM) within the same neurons, we designed a WM-guided feature attention task requiring to maintain a visual feature in WM, and subsequently use that feature to guide attention. Animals were required to fixate on a small yellow square (0.6-by-0.6 visual degrees) presented at the center of the screen during the entirety of each task trial (fixation window radius: 2.6 visual degrees). Eye movements away from this window terminated a trial without reward. Monkeys began each trial (Fig. 1A) by grasping a lever. After a 1000 ms fixation period, a cue stimulus was presented for 800 ms, consisting of a full-screen surface of random dots with coherent motion (5 dots/deg^2^; 0.15 deg dot diameter; 10.9deg/s dot speed, 667 ms dot life) in one of two possible directions (45° clockwise from upward or 45° clockwise from downward). Monkey Sh was further trained to perform the task with four possible cue directions separated by 90°. After a 3200 ms delay period containing the fixation square alone, two test stimuli were presented, consisting of two overlapping full-screen random-dot surfaces with the same parameters as the cue; one of them – the target surface – matched the cue motion direction, whereas the other – the distractor surface – had motion in the opposite direction. Subsequently, a squared patch of dots (length 8.14 degrees) at a random location within the target surface increased motion speed (to double the original) for 250 ms and returned to its initial speed. The target speed change onset occurred at a random time between 900 and 2700 ms from test onset. In half of the trials, the speed-changing patch occurred in the distractor surface, followed by a second patch occurring in the target surface at a random time (minimum 800 ms after the distractor patch and maximum 2700 ms from test onset). Monkeys were required to selectively attend to the test surface matching the cue direction (i.e., the target) in order to detect the occurrence of the target speed change and report it by releasing the lever, while ignoring distractor changes. A correct trial occurred when monkeys released the lever within a response window of 120 to 580 ms from target change onset; a 1 mL juice reward was then delivered to the monkey with an automated juicer. The reaction time was measured as the time between target onset and lever release. Lever releases during the test period (after 900 ms) but before the target were classified as early response errors; releases within 120 to 580 ms from distractor onset were classified as distractor response errors. Trials with no lever release before the end of the response window were classified as no-response errors. Task performance was quantified as the percentage of correct trials over the sum of all correct and error trials defined above. Trials with either a lever release before the test or a fixation loss were excluded from the analyses.

We varied the perceptual difficulty of the task between trials by varying the percentage of speed-changing dots within the target/distractor patches (85%, easier; 55%, intermediate; or 35%, difficult). These trials were presented randomly and with equal frequency. Difficult trials imposed higher attentional demands to correctly perform the task, thus ensuring that monkeys selectively attended the target surface throughout the session. We confirmed this by ensuring that performance in difficult trials was considerably higher than expected by chance in all sessions. However, only including difficult trials would risk monkeys losing motivation to perform the task or eventually un-learning the task rule due to low performance. This was prevented by including easier trials. In a pilot test, naïve human subjects instructed to perform the task were generally unable to detect most speed changes until they had trained for several trials to selectively attend to the target while ignoring the distractor.

### Surgical procedures

#### Head post and cranial chamber implantation

All surgical procedures were performed while animals were under anesthesia. Animals were provided analgesics and antibiotics after surgery. Each monkey was first surgically implanted with a titanium headpost over the posterior end of the skull. After recovery was complete, animals began task training while being head-fixed with a head-post holder. After the period of training was completed, a second surgery was performed for implantation of cranial chambers. We first performed round craniotomies over left and right prefrontal cortex, and left parietal cortex. We then performed a durotomy in each prefrontal cranial window and implanted a round transparent silicone artificial dura so that its edge was slid under the native dura around the perimeter of the durotomy. Subsequently, we implanted three cranial chambers over the three craniotomies. Each prefrontal chamber was designed so that its bottom edge was inserted into the craniotomy and touched the artificial dura (instead of sitting on the skull surface like standard implants). The transparent artificial dura windows allowed for multiple optogenetic and electrophysiological procedures including virus injections, continuous *in vivo* visual tracking of viral expression, minimally-invasive external optical stimulation, and electrophysiological recordings.

Chambers were made of polyether ether ketone (PEEK) and had an inner diameter of 19 mm. Using structural MR images of the skull and brain, each chamber was custom-designed with a base to fit the surrounding skull (Mulliken et al., 2015), and was attached to it with ceramic screws. The headpost implanted in the first surgery was similarly custom-fit to the skull and attached to it with titanium screws. All custom-fit implants were fabricated with a 5-axis CNC machine. The parietal chamber implant had a cylinder protruding over the base, onto which the electrophysiological recording towers were mounted. Prefrontal chamber implants were designed without a protruding cylinder; instead, the cylinder was screwed and affixed to the base before recording sessions. The three chambers were placed over left and right prefrontal cortex, and left parietal cortex. Prefrontal chambers were positioned to include the region between the arcuate sulcus and the posterior end of the principal sulcus. The parietal chamber was placed and oriented for probe trajectories to access areas LIP, MST, and MT as perpendicularly as possible to the cortical sheet (Fig. 1C).

Chambers were cleaned 3 to 5 times per week under aseptic conditions. The prefrontal chambers were rinsed using a sterile saline antibiotic cocktail with 2.5% Penicillin G Sodium powder (USP 5 million units) and 5% Amikacin (0.25 g/mL). After rinsing, a gauze piece was placed on the artificial dura with a drop of a second saline cocktail (16% Penicillin and 40% Amikacin); a silicone disc was then placed over the gauze to maintain moderate pressure on the dura, and a water-tight cap was placed and screwed on top of the chamber. The parietal chamber was rinsed using sterile saline and betadine solution. During chamber cleaning, we also monitored the health of the prefrontal cortical tissue, as well as the progression of native dura regrowth. When enough dura had grown to occlude visibility of the cortical surface, we repeated the durotomy procedure following all surgical standards described above. This was only necessary after 6 months from the initial durotomy, and only once for each monkey during the period of experimental sessions.

A structural magnetic resonance imaging (MRI) scan was performed after the implantation surgery to obtain an anatomical map of the chamber locations. Each chamber’s recording grid (see Electrophysiological recordings section) was filled with a 1% agar solution in sterile water with 0.8% Ablavar (gadolinium-based contrast agent) and placed inside the chamber for the duration of the scan. The MR images served to map the position of the chamber and grid, and the transcortical probe trajectories corresponding to all grid slots (Fig. 1C).

#### Virus injections

To induce neuronal expression of the opsin Jaws in left and right LPFC-p, we performed injections of the AAV5.hSyn.Jaws-KGC.GFP.ER2-WPRE.hG viral construct. The following setup was prepared identically for each prefrontal hemisphere. Injections were made with a 9.6 cm long 31 GA needle. The needle was reinforced by inserting it and gluing it to an 8.3 cm-long 23 GA needle, keeping 8mm of the tip exposed for injection penetration. The reinforcement needle was held by a microdrive tower (NAN Instruments, Israel) on an XY table fixed to the chamber. The needle was attached to a 70-cm Intramedic polyelthylene tubing, which was in turn attached to a 25-μL Hamilton syringe mounted on a syringe pump (Harvard Apparatus, USA). These attachments were sealed with Bondic glue and made air-tight.

Virus injections were performed two to three weeks after chamber implantation, under the same surgical conditions described above. The left and right prefrontal chambers were first cleaned. Then, the towers with injection needles were mounted on the chambers. In each hemisphere, we selected twelve injection locations in a grid-like layout distributed across the LPFC-p subregion between the arcuate sulcus and the posterior end of the principal sulcus. Neighboring injection locations were approximately 1.6 mm apart. Locations were shifted away from any visible blood vessel. Injections were performed serially at each of the twelve locations, beginning with the most central location and spiraling outwards along the grid of planned locations. The needle was quickly shifted between injection locations using the microdrive tower’s XY table. At each location, we lowered the needle to penetrate the artificial dura and cortex, and we stopped at a depth of 4 mm below where the needle had dimpled the dura. We performed the first injection at that depth (1 μL), and subsequent injections by retracting the needle in steps of 1 mm until 2 mm deep (1 μL), and then in steps of 0.5 mm (0.5 μL) until the needle was out of the cortex.

The above surgical implantation and injection procedures were developed and tested on a pilot monkey prior to performing them on the two trained monkeys. This allowed the materials and procedures to be optimized. In one prefrontal chamber of the pilot monkey, we injected three different versions of the Jaws viral construct: one with AAV5, and two with AAV8, from different vector cores/batches. Visual inspection of GFP epifluorescence through the artificial dura determined that the AAV5 construct led to the most widespread expression. This determined the construct selection for the experiments.

### Electrophysiological recordings

We recorded intracortical neuronal signals using V-Probes and S-Probes (Plexon Inc., USA) – multi-contact linear electrophysiological probes (i.e., laminar probes). Probes were 110 to 130 mm long and had 16 or 32 contacts (i.e., channels) with 150 or 100 μm spacing, respectively. Each probe was mounted on an electric microdrive tower (NAN Instruments, Israel) attached to the chamber, and was embedded in a guide tube (costume-cut spinal needle; sharp for parietal probes, blunt for prefrontal probes). The probe and guide tube were moved independently of each other by an electric microdrive and by hand, respectively. In preparation for a recording session, we mounted one probe on each prefrontal chamber and three on the parietal chamber. A plastic grid was placed in the parietal chamber, and the guide tube with the probe was placed inside a grid slot. Using the structural MRI images, the grid slot coordinates of the probe and its planned final depth were selected to target the cortical location of interest (Fig. 1C).

After mounting all microdrive towers on the chambers, we manually lowered all guide tubes to poke through the native dura (for parietal chamber recordings) or to press and dimple the artificial dura (for prefrontal chamber recordings). The probes were then slowly inserted into the cortex using the computer-controlled electric microdrive, stopping at the planned final depth. The exact target depth was adjusted based on the presence of single- and multi-unit spiking activity across a large range of recording channels. In each session, we recorded activity from four to five probes simultaneously – one per prefrontal chamber and three in the parietal chamber. The three parietal chamber probes were placed to target areas LIP, MST and MT.

In each area, we targeted neurons with motion direction selectivity during at least one of the task periods (cue, delay or test). Probe recordings with no direction-selective neurons were excluded from analysis, and the location was excluded from subsequent recording sessions. Recording locations were selected to be more proximal to previous recording locations with direction selective neurons and more distal from any locations without these neurons.

### Optogenetic stimulation

Optogenetic stimulation was performed with two red lasers (635 nm) with adjustable power up to 4W (Shanghai Laser & Optics Century Co., Ltd., Shanghai). Each laser was coupled with a 2-meter long step index optical fiber with a ceramic ferrule ending (400 μm core diameter, 2.5 mm optical density). Each fiber was firmly attached to a microdrive tower (parallel to it) fixed to the chamber so that the fiber ending was located above the LPFC-p cortical surface on each hemisphere, pointing towards the center of the opsin expression region. During recording sessions, the LPFC-p chamber was sealed with two layers of black electrical tape that prevented the laser light from escaping the chamber. The distance of the fiber ending to the cortical surface ranged between 17 and 29 mm, and was chosen so the laser beam would cover the entire region of opsin expression. The laser beam diameter ranged between 8.5 and 9.5 mm in monkey Sh, and between 10 and 14.5 mm in monkey St. The laser power was adjusted so that the power density fell within the range of 30 to 42 mW/mm^2^. In pilot experiments with a test monkey, this power range was found to successfully yield optogenetic inactivation of neurons across all cortical depths. Importantly, a previous study showed that cortical stimulation with a 635 nm laser at 100 mW/mm^2^ (more than twice the power density used here) did not result in tissue heating greater than 1°C (Acker et al., 2016).

In each recording session, the two lasers were simultaneously turned on in specific task periods in half of the trials (opto trials) and not in the remaining half (control trials). Opto and control trials were randomly interleaved. The lasers were automatically controlled by the behavioral task computer via TTL pulses. In a fraction of the sessions (delay-opto), the lasers were turned on for 2 s during the delay period, starting 300 ms after the sample stimulus offset, and ending 900 ms before the test onset. In the remaining sessions (test-opto), the lasers were turned on at the test onset, lasting for 2 s or until the monkey made a response. The laser stimulation regime was a single square pulse. Delay-opto and test-opto sessions were randomly interleaved, with sessions usually separated by at least a day in between.

### Data analysis

Automatic spike detection and spike sorting were performed on the electrophysiological signals recorded from each channel in each session using Spyking Circus (Yger et al., 2018). This yielded the spike timestamps of single neurons during task trials at 1 ms sampling rate. All subsequent analyses were performed on Matlab (Mathworks, Inc.). For each neuron, we calculated the firing rate over time across each trial in 50 ms bins. Trials were grouped by each combination of cue direction condition and optogenetic stimulation condition (control, delay-opto, test-opto). Using a Matlab function (ROC; Cardillo G. (2008) ROC curve: compute a Receiver Operating Characteristics curve. http://www.mathworks.com/matlabcentral/fileexchange/19950), we performed Receiver Operating Characteristics analysis to obtain the area under the ROC curve (auROC) comparing the distributions of firing rates across trials in pairs of conditions with opposite cue directions. This was done independently on the firing rate in the cue period (1000 to 1800 ms from trial onset), delay period (2100 to 4100 ms), and test period (5000 to 5900 ms), and independently on control trials and opto trials. For each neuron, this analysis yielded the magnitude and significance of sensory discriminability, WM discriminability and attentional discriminability in control and opto trials. Neurons with significant direction discriminability in at least one of the three periods (cue, delay or test) were included for subsequent analyses. Given our aim to measure the effects of optogenetic LPFC-p inactivation on task performance and on each neuron’s WM coding and feature attentional modulation strength, our analyses included correct and error trials (target responses, early responses, distractor responses and no responses). As described previously, monkey Sh was trained to perform the task with 4 possible cue directions. For this monkey, ROC analysis in each task period was performed on each pair of opposite cue direction conditions. If one pair of directions showed significant auROC, this pair was chosen for subsequent analyses. If both direction pairs showed significant auROC, the pair yielding the highest discriminability (likely the most aligned to the neuron’s direction preference axis) was selected for further analyses. From the selected pair of directions, the one for which the neuron showed the highest mean activity across trials was identified as the preferred direction.

To examine whether test period LPFC-p inactivation caused a reduction in attention discriminability in each area (Fig. 5J), we tested whether the mean attention auROC across all neurons with significant attention discriminability in test opto trials was significantly lower than the mean attention auROC across all neurons with significant attention discriminability in control trials, using a one-tailed Wilcoxon rank sum test. A comparable procedure was performed on the WM auROC values to examine whether delay period LPFC-p inactivation caused a reduction in WM discriminability in each area (Fig 6H), and on the attention auROC values to examine whether delay period LPFC-p inactivation caused a reduction in attention discriminability in each area (Fig 6K).

In the population analyses used to produce the results shown in Fig. 4E-K, 5B-E, 6B-D, and S1A-H, the firing rate over time of each neuron in each condition was normalized by dividing by its mean firing rate across the entire time period of interest and across all conditions compared. Plots of mean firing rate over time (Fig. 1,2,4,5,6, S1) were obtained by convolving the binned firing rates in each trial with a 200 ms kernel and then obtaining the mean and standard error of these traces across trials. This data was then used to calculate auROC over time for each neuron and obtain the mean auROC and standard error across neurons over time (Fig. 5F-I, 6E-G). For each neuron, the auROC expected by chance was obtained by repeating the same procedure after randomly permuting the control and opto condition labels between trials. We tested for inactivation-induced reductions in attention or WM discriminability over time in each area by comparing the mean auROC values across neurons between opto and control conditions at all test or delay period time bins using one-tailed Wilcoxon rank sum tests corrected for multiple comparisons.

## Acknowledgements

We thank Hang Le, Briana McRae, and Callie Kunz with assistance with spike sorting; Rick Ono for assistance with anatomical mapping of recorded brain regions; Amanda Marino and Tal Inbar for assistance with behavioral training, implant care, and probe implantation; John Reynolds and Anirvan Nandy for guidance with artificial dura implant methods. This work was supported by NIH R01-EY029666.

## Author Contributions

Funding acquisition: RD

Study design: DMH, RD

Implant design, surgical procedures, implant care: DMH, HX

Animal training: DMH

Electrophysiology data collection and analysis: DMH

Probe implantation: DMH, FACA

Writing, first draft: DMH

Writing, revised draft: DMH, RD, HX

## Supplementary Figures

**Figure S1.**
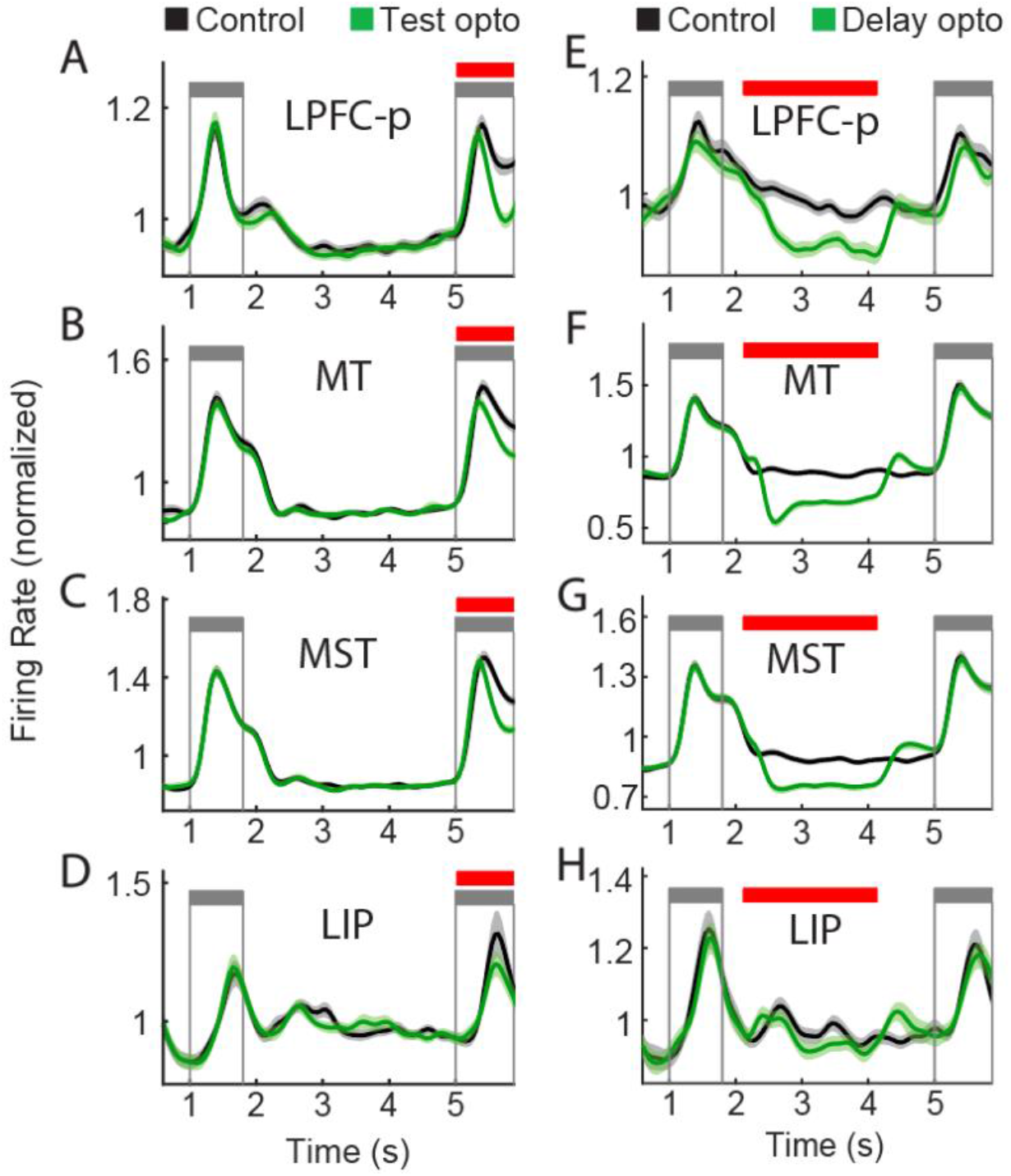
Effects of test and delay period optogenetic prefrontal inactivation on firing rates across all neurons in each area. Mean normalized firing rate (± standard error) over time in control and opto trials across all neurons in LPFC, MT, MST, and LIP during sessions with optogenetic stimulation during the test period (A-D) or delay period (E-H). Each neuron’s firing rate was normalized by its mean firing rate across all trial periods and conditions. Firing rate decreases are visible in the mean across all neurons, although less pronounced that in the mean across significantly modulated neurons (Fig. 4).

**Figure S2.**
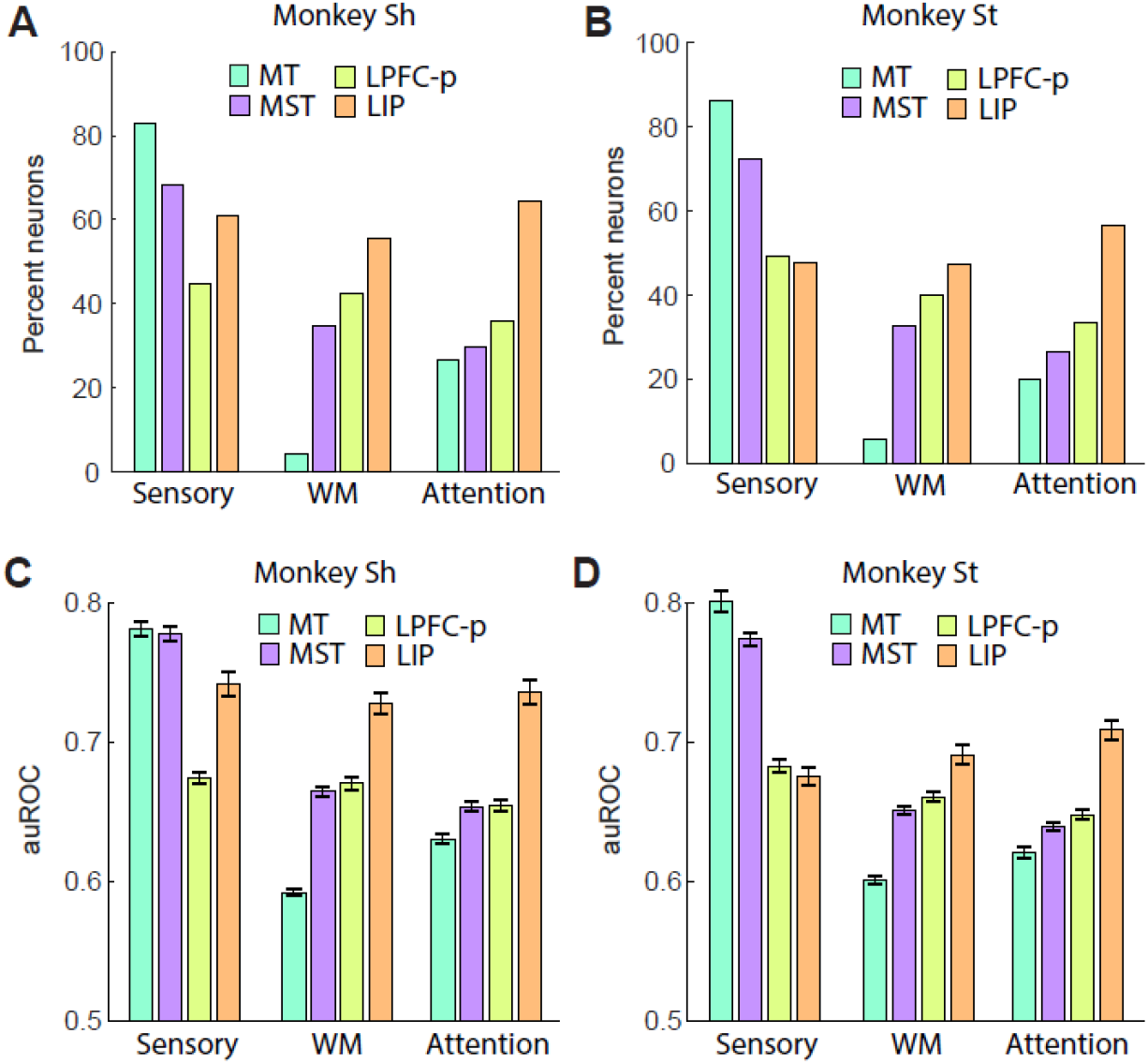
Effects of WM and attention shown separately for the two monkeys. (A,B) Percentage of neurons with significant sensory, WM, and attentional discriminability in each brain area, shown separately for the two monkeys. (C,D) Mean (± standard error) sensory, WM, and attentional discriminability across all neurons in each brain area, shown separately for the two monkeys.

## Notes

### Competing Interest Statement

The authors have declared no competing interest.

## References

Acker, L., Pino, E. N., Boyden, E. S., & Desimone, R. (2016). FEF inactivation with improved optogenetic methods. Proceedings of the National Academy of Sciences of the United States of America, 113(46), E7297–E7306. https://doi.org/10.1073/PNAS.1610784113/SUPPL_FILE/PNAS.1610784113.SAPP.PDF

Awh, E., & Jonides, J. (2001). Overlapping mechanisms of attention and spatial working memory. Trends in Cognitive Sciences, 5(3), 119–126. https://doi.org/10.1016/S1364-6613(00)01593-X

Bae, G. Y., & Luck, S. J. (2018). Dissociable Decoding of Spatial Attention and Working Memory from EEG Oscillations and Sustained Potentials. Journal of Neuroscience, 38(2), 409–422. https://doi.org/10.1523/JNEUROSCI.2860-17.2017

Bichot, N. P., Heard, M. T., DeGennaro, E. M., & Desimone, R. (2015). A Source for Feature-Based Attention in the Prefrontal Cortex. Neuron, 88(4), 832–844. https://doi.org/10.1016/J.NEURON.2015.10.001

Bichot, N. P., Rossi, A. F., & Desimone, R. (2005). Parallel and serial neural mechanisms for visual search in macaque area V4. Science (New York, N.Y.), 308(5721), 529–534. https://doi.org/10.1126/SCIENCE.1109676

Bichot, N. P., Xu, R., Ghadooshahy, A., Williams, M. L., & Desimone, R. (2019). The role of prefrontal cortex in the control of feature attention in area V4. Nature Communications 2019 10:1, 10(1), 1–12. https://doi.org/10.1038/s41467-019-13761-7

Buschman, T. J., & Miller, E. K. (2007). Top-down versus bottom-up control of attention in the prefrontal and posterior parietal cortices. Science, 315(5820), 1860–1864. https://doi.org/10.1126/SCIENCE.1138071/SUPPL_FILE/BUSCHMAN.SOM.PDF

Christophel, T. B., Klink, P. C., Spitzer, B., Roelfsema, P. R., & Haynes, J. D. (2017). The Distributed Nature of Working Memory. Trends in Cognitive Sciences, 21(2), 111–124. https://doi.org/10.1016/J.TICS.2016.12.007

Chuong, A. S., Miri, M. L., Busskamp, V., Matthews, G. A. C., Acker, L. C., Sørensen, A. T., Young, A., Klapoetke, N. C., Henninger, M. A., Kodandaramaiah, S. B., Ogawa, M., Ramanlal, S. B., Bandler, R. C., Allen, B. D., Forest, C. R., Chow, B. Y., Han, X., Lin, Y., Tye, K. M., … Boyden, E. S. (2014). Noninvasive optical inhibition with a red-shifted microbial rhodopsin. Nature Neuroscience 2014 17:8, 17(8), 1123–1129. https://doi.org/10.1038/nn.3752

Collins CE, Airey DC, Young NA, Leitch DB, & Kaas JH. (2010) Neuron densities vary across and within cortical areas in primates. Proceedings of the National Academy of Sciences, 107(36):15927–15932. https://doi.org/10.1073/pnas.1010356107

Desimone, R., & Duncan, J. (1995). Neural mechanisms of selective visual attention. Annual Review of Neuroscience, 18, 193–222. https://doi.org/10.1146/ANNUREV.NE.18.030195.001205

Fenno, L., Yizhar, O., & Deisseroth, K. (2011). The Development and Application of Optogenetics. https://Doi.Org/10.1146/Annurev-Neuro-061010-113817, 34, 389–412. https://doi.org/10.1146/ANNUREV-NEURO-061010-113817

Gong, X., Mendoza-Halliday, D., Ting, J. T., Kaiser, T., Sun, X., Bastos, A. M., Wimmer, R. D., Guo, B., Chen, Q., Zhou, Y., Pruner, M., Wu, C. W. H., Park, D., Deisseroth, K., Barak, B., Boyden, E. S., Miller, E. K., Halassa, M. M., Fu, Z., … Feng, G. (2020). An Ultra-Sensitive Step-Function Opsin for Minimally Invasive Optogenetic Stimulation in Mice and Macaques. Neuron, 107(1), 38-51.e8. https://doi.org/10.1016/J.NEURON.2020.03.032

Gregoriou, G. G., Rossi, A. F., Ungerleider, L. G., & Desimone, R. (2014). Lesions of prefrontal cortex reduce attentional modulation of neuronal responses and synchrony in V4. Nature Neuroscience 2014 17:7, 17(7), 1003–1011. https://doi.org/10.1038/nn.3742

Harris, A. M., Jacoby, O., Remington, R. W., Becker, S. I., & Mattingley, J. B. (2020). Behavioral and electrophysiological evidence for a dissociation between working memory capacity and feature-based attention. Cortex, 129, 158–174. https://doi.org/10.1016/J.CORTEX.2020.04.009

Ibos, G., & Freedman, D. J. (2016). Interaction between Spatial and Feature Attention in Posterior Parietal Cortex. Neuron, 91(4), 931–943. https://doi.org/10.1016/J.NEURON.2016.07.025

Jonikaitis, D., & Moore, T. (2019). The interdependence of attention, working memory and gaze control: behavior and neural circuitry. Current Opinion in Psychology, 29, 126–134. https://doi.org/10.1016/J.COPSYC.2019.01.012

Konecky, R. O., Smith, M. A., & Olson, C. R. (2017). Monkey prefrontal neurons during sternberg task performance: Full contents of working memory or most recent item? Journal of Neurophysiology, 117(6), 2269–2281. https://doi.org/10.1152/JN.00541.2016/ASSET/IMAGES/LARGE/Z9K0061741380008.JPEG

Lara, A. H., & Wallis, J. D. (2014). Executive control processes underlying multi-item working memory. Nature Neuroscience 2014 17:6, 17(6), 876–883. https://doi.org/10.1038/nn.3702

Leavitt, M. L., Mendoza-Halliday, D., & Martinez-Trujillo, J. C. (2017). Sustained Activity Encoding Working Memories: Not Fully Distributed. Trends in Neurosciences, 40(6), 328–346. https://doi.org/10.1016/J.TINS.2017.04.004

Lebedev, M. A., Messinger, A., Kralik, J. D., & Wise, S. P. (2004). Representation of Attended Versus Remembered Locations in Prefrontal Cortex. PLOS Biology, 2(11), e365. https://doi.org/10.1371/JOURNAL.PBIO.0020365

Mattfeld, A. T., Whitfield-Gabrieli, S., Biederman, J., Spencer, T., Brown, A., Fried, R., & Gabrieli, J. D. E. (2016). Dissociation of working memory impairments and attention-deficit/hyperactivity disorder in the brain. NeuroImage: Clinical, 10, 274–282. https://doi.org/10.1016/J.NICL.2015.12.003

Maunsell, J. H. R., & Treue, S. (2006). Feature-based attention in visual cortex. Trends in Neurosciences, 29(6), 317–322. https://doi.org/10.1016/J.TINS.2006.04.001

Mayer, J. S., Roebroeck, A., Maurer, K., & Linden, D. E. J. (2010). Specialization in the default mode: Task-induced brain deactivations dissociate between visual working memory and attention. Human Brain Mapping, 31(1), 126. https://doi.org/10.1002/HBM.20850

McAdams, C. J., & Maunsell, J. H. R. (1999). Effects of Attention on Orientation-Tuning Functions of Single Neurons in Macaque Cortical Area V4. Journal of Neuroscience, 19(1), 431–441. https://doi.org/10.1523/JNEUROSCI.19-01-00431.1999

Mendoza-Halliday, D., & Martinez-Trujillo, J. C. (2017). Neuronal population coding of perceived and memorized visual features in the lateral prefrontal cortex. Nature Communications 2017 8:1, 8(1), 1–13. https://doi.org/10.1038/ncomms15471

Mendoza-Halliday, D., Torres, S., & Martinez-Trujillo, J. C. (2014). Sharp emergence of feature-selective sustained activity along the dorsal visual pathway. Nature Neuroscience 2014 17:9, 17(9), 1255–1262. https://doi.org/10.1038/nn.3785

Messinger, A., Lebedev, M. A., Kralik, J. D., & Wise, S. P. (2009). Multitasking of Attention and Memory Functions in the Primate Prefrontal Cortex. Journal of Neuroscience, 29(17), 5640–5653. https://doi.org/10.1523/JNEUROSCI.3857-08.2009

Mongillo, G., Barak, O., & Tsodyks, M. (2008). SynaptiC Theory of Working Memory. Science, 319(5869), 1543–1546. https://doi.org/10.1126/SCIENCE.1150769/SUPPL_FILE/MONGILLO.SOM.PDF

Mulliken, G. H., Bichot, N. P., Ghadooshahy, A., Sharma, J., Kornblith, S., Philcock, M., & Desimone, R. (2015). Custom-fit radiolucent cranial implants for neurophysiological recording and stimulation. Journal of Neuroscience Methods, 241, 146–154. https://doi.org/10.1016/J.JNEUMETH.2014.12.011

Nandy, A., Nassi, J. J., Jadi, M. P., & Reynolds, J. (2019). Optogenetically induced low-frequency correlations impair perception. ELife, 8. https://doi.org/10.7554/ELIFE.35123

Nandy, A. S., Nassi, J. J., & Reynolds, J. H. (2017). Laminar Organization of Attentional Modulation in Macaque Visual Area V4. Neuron, 93(1), 235–246. https://doi.org/10.1016/J.NEURON.2016.11.029

Nassi, J. J., Avery, M. C., Cetin, A. H., Roe, A. W., & Reynolds, J. H. (2015). Optogenetic Activation of Normalization in Alert Macaque Visual Cortex. Neuron, 86(6), 1504–1517. https://doi.org/10.1016/J.NEURON.2015.05.040

Olivers, C. N. L. (2008). Interactions between visual working memory and visual attention. Frontiers in Bioscience : A Journal and Virtual Library, 13(3), 1182–1191. https://doi.org/10.2741/2754

Paxinos, G. (2009). The rhesus monkey brain in stereotaxic coordinates. 410. https://books.google.com/books/about/The_Rhesus_Monkey_Brain_in_Stereotaxic_C.html?id=7HW6HgAACAAJ

Peers, P. v., Astle, D. E., Duncan, J., Murphy, F. C., Hampshire, A., Das, T., & Manly, T. (2020). Dissociable effects of attention vs working memory training on cognitive performance and everyday functioning following fronto-parietal strokes. Neuropsychological Rehabilitation, 30(6), 1092–1114. https://doi.org/10.1080/09602011.2018.1554534/SUPPL_FILE/PNRH_A_1554534_SM9365.JPG

Petrides, M. (1991). Monitoring of selections of visual stimuli and the primate frontal cortex. Proceedings. Biological Sciences, 246(1317), 293–298. https://doi.org/10.1098/RSPB.1991.0157

Petrides, M. (2005). Lateral prefrontal cortex: architectonic and functional organization. Philosophical Transactions of the Royal Society B: Biological Sciences, 360(1456), 781–795. https://doi.org/10.1098/RSTB.2005.1631

Rossi, A. F., Pessoa, L., Desimone, R., & Ungerleider, L. G. (2009). The prefrontal cortex and the executive control of attention. Experimental Brain Research, 192(3), 489–497. https://doi.org/10.1007/S00221-008-1642-Z/METRICS

Ruiz, O., Lustig, B. R., Nassi, J. J., Cetin, A., Reynolds, J. H., Albright, T. D., Callaway, E. M., Stoner, G. R., & Roe, A. W. (2013). Optogenetics through windows on the brain in the nonhuman primate. Journal of Neurophysiology, 110(6), 1455–1467. https://doi.org/10.1152/JN.00153.2013

Stokes, M. G. (2015). ‘Activity-silent’ working memory in prefrontal cortex: a dynamic coding framework. Trends in Cognitive Sciences, 19(7), 394–405. https://doi.org/10.1016/J.TICS.2015.05.004

Tas, A. C., Luck, S. J., & Hollingworth, A. (2016). The Relationship between Visual Attention and Visual Working Memory Encoding: A Dissociation between Covert and Overt Orienting. Journal of Experimental Psychology. Human Perception and Performance, 42(8), 1121. https://doi.org/10.1037/XHP0000212

Tomasi, D., Chang, L., Caparelli, E. C., & Ernst, T. (2007). Different activation patterns for working memory load and visual attention load. Brain Research, 1132(1), 158–165. https://doi.org/10.1016/J.BRAINRES.2006.11.030

Tremblay, S., Acker, L., Afraz, A., Albaugh, D. L., Amita, H., Andrei, A. R., Angelucci, A., Aschner, A., Balan, P. F., Basso, M. A., Benvenuti, G., Bohlen, M. O., Caiola, M. J., Calcedo, R., Cavanaugh, J., Chen, Y., Chen, S., Chernov, M. M., Clark, A. M., … Platt, M. L. (2020). An Open Resource for Non-human Primate Optogenetics. Neuron, 108(6), 1075-1090.e6. https://doi.org/10.1016/J.NEURON.2020.09.027

Tremblay, S., Pieper, F., Sachs, A., & Martinez-Trujillo, J. (2015). Attentional filtering of visual information by neuronal ensembles in the primate lateral prefrontal cortex. Neuron, 85(1), 202–215. https://doi.org/10.1016/J.NEURON.2014.11.021

Treue, S., & Martínez Trujillo, J. C. (1999). Feature-based attention influences motion processing gain in macaque visual cortex. Nature 1999 399:6736, 399(6736), 575–579. https://doi.org/10.1038/21176

Warden, M. R., & Miller, E. K. (2010). Task-Dependent Changes in Short-Term Memory in the Prefrontal Cortex. Journal of Neuroscience, 30(47), 15801–15810. https://doi.org/10.1523/JNEUROSCI.1569-10.2010

Xu, R., Bichot, N. P., Takahashi, A., & Desimone, R. (2022). The cortical connectome of primate lateral prefrontal cortex. Neuron, 110(2), 312-327.e7. https://doi.org/10.1016/J.NEURON.2021.10.018

Yger P., Spampinato, G.L.B, Esposito E., Lefebvre B., Deny S., Gardella C., Stimberg M., Jetter F., Zeck G. Picaud S., Duebel J., Marre O., A spike sorting toolbox for up to thousands of electrodes validated with ground truth recordings in vitro and in vivo, eLife 2018;7:e34518. https://doi.org/10.7554/eLife.34518

Young NA, Collins CE, & Kaas JH. (2013) Cell and neuron densities in the primary motor cortex of primates. Front. Neural Circuits, 7:30. https://doi.org/10.3389/fncir.2013.00030

Zaksas, D., & Pasternak, T. (2006). Directional signals in the prefrontal cortex and in area MT during a working memory for visual motion task. The Journal of Neuroscience : The Official Journal of the Society for Neuroscience, 26(45), 11726–11742. https://doi.org/10.1523/JNEUROSCI.3420-06.2006

